# Metabolomics profiling reveals distinct, sex-specific signatures in the serum and brain metabolomes in the mouse models of Alzheimer’s disease

**DOI:** 10.1101/2023.12.22.573059

**Authors:** Ravi S. Pandey, Mattias Arnold, Richa Batra, Jan Krumsiek, Kevin P. Kotredes, Dylan Garceau, Harriet Williams, Michael Sasner, Gareth R. Howell, Rima Kaddurah-Daouk, Gregory W. Carter

## Abstract

**INTRODUCTION:** Increasing evidence suggests that metabolic impairments contribute to early Alzheimer’s disease (AD) mechanisms and subsequent dementia. Signals in metabolic pathways conserved across species provides a promising entry point for translation.

**METHODS:** We investigated differences of serum and brain metabolites between the early-onset 5XFAD and late-onset LOAD1 (APOE4.Trem2*R47H) mouse models of AD to C57BL/6J controls at six months of age.

**RESULTS:** We identified sex differences for several classes of metabolites, such as glycerophospholipids, sphingolipids, and amino acids. Metabolic signatures were notably different between brain and serum in both mouse models. The 5XFAD mice exhibited stronger differences in brain metabolites, whereas LOAD1 mice showed more pronounced differences in serum.

**DISCUSSION:** Several of our findings were consistent with results in humans, showing glycerophospholipids reduction in serum of APOE4 carriers and replicating the serum metabolic imprint of the APOE4 genotype. Our work thus represents a significant step towards translating metabolic dysregulation from model organisms to human AD.

## INTRODUCTION

Alzheimer’s disease (AD) is a devastating neurodegenerative disorder and is the leading cause of dementia. Neuropathologically, AD is characterized by the accumulation of amyloid plaques and tau fibrillary tangles in the brain [1-3]. Based on the age of onset AD is characterized into early and late onset AD. Early-onset Alzheimer’s disease (EOAD, also known as familial AD or FAD) strikes prior to the age of 65 and mutations in genes coding for *amyloid precursor protein (APP), presenilin 1 (PSEN1)*, and *presenilin 2 (PSEN2)* were identified in familial AD cases [4, 5]. However, sporadic or late-onset AD (LOAD) is the most common form of AD (>95% prevalence), in which symptoms arise after 65 years of age [5, 6] and its risk factors are age, the apolipoprotein E ε4 allele (APOE ε4) and rare point mutations in the triggering receptor expressed on myeloid cells 2 (TREM2) [3, 7-10]. In addition, females are at higher risk for AD [11, 12] and approximately 65% of Americans with AD are females [13]. In addition, female *APOE* ε4 carriers are at greater risk of developing AD compared to men with the same variant [14-16].

Pathophysiological changes associated with AD begin decades before the manifestation of clinical symptoms [11]. Metabolic decline is one of the earliest symptoms detected, with reduced glucose utilization using fluorodeoxyglucose positron emission tomography (FDG-PET) in patients with mild cognitive impairment (MCI), which can be an early stage of AD [17]. Furthermore, disruption in glucose metabolism associated with early mitochondrial dysfunction is detected in multiple animal models and AD-affected individuals[18]. Perturbations in multiple metabolic networks such as lysine metabolism, tricarboxylic acid (TCA) cycle, and lipid metabolism were reported in MCI individuals compared to healthy individuals [19]. This suggests that metabolic dysfunction could play an important role in early stages of AD.

Metabolic dysregulation one of the hallmarks of AD impact of all three of the major AD risk factors: age, *APOE* genotype in European cohorts, and sex [20-23]. A recent study in the Alzheimer’s Disease Neuroimaging Initiative (ADNI) cohorts identified the effect of sex and APOE *ε*4 status on metabolic alternations related to AD biomarkers [12]. Even in mouse models of AD, changes in metabolic pathways related to energetic stress were more pronounced in female mice compared to males [19, 24]. However, the molecular mechanisms underlying these sex-linked differences remain to be elucidated.

Heterogeneity in humans due to genetic diversity and the impact of environmental and lifestyle differences complicates the molecular study of disease mechanisms [25]. Additionally, for human studies, experimental design is often limited by sample availability, reducing confidence in inferences [11]. Animal models have been critical for understanding the development and progression of AD and offer opportunities to study the effects of disease related risk factors in a controlled environment [3, 25]. Genetic mouse models of AD further enable the controlled collection of cross-sectional sampling at multiple ages and analysis of metabolic changes during various disease stages. Signals in conserved metabolic pathways could thus provide a method to translate experimental findings in preclinical mouse models to humans [11]. Transgenic mouse models with gene mutations for APP and PSEN1 were widely used to investigate the biofluids and brain metabolome and observed significant overlap in the affected metabolic pathways identified in AD patients [26-28]. However, these mouse models were limited to fAD transgenic models that represent a small number of AD cases, and previous studies did not interrogate the influence of sex-specific differences in metabolic changes.

To fill this gap, we comprehensively profiled the serum and brain metabolomes of APOE4.Trem2*R47H mice (a genetic model for LOAD), the 5XFAD mice (a transgenic amyloid model), and C57BL/6J (control) mice at six months of age in both sexes. A total of 142 metabolites were measured in both brain and serum, including phospholipids, amino acids, biogenic amines, and acylcarnitines. The LOAD1 double mutant strain carries two primary risk alleles, humanized APOE4 and the Trem2*R47H variant on the C57BL/6J (B6) background [29]. The 5XFAD transgenic mice overexpress human APP with three FAD mutations and human PSEN1 with two FAD mutations on the B6 background [30]. We investigated the sex differences in metabolic effects for the 5XFAD and the APOE4 genotype in both blood and serum metabolomes as well as the correspondence between brain and serum metabolite levels in mouse models. We further compared our findings with recent results from the Alzheimer’s Disease Neuroimaging Initiative (ADNI) cohorts [12], as well as human serum and brain metabolomics data from the Rush Religious Order Study and Memory and Aging Project (ROS/MAP) [31]. All datasets described in this study are available to the research community through the AD Knowledge Portal (https://adknowledgeportal.synapse.org/).

## METHODS AND MATERIALS

### Animal models

All animal models were obtained from The Jackson Laboratory. All experiments were approved by the Animal Care and Use Committee at The Jackson Laboratory. LOAD1 mice (JAX #28709) carry a humanized version of the prominent *APOEε4* genetic risk factor for LOAD (late-onset Alzheimer’s disease), and a relatively rare deleterious variant R47H allele of *Trem2* gene [29], while the 5XFAD transgenic mice (JAX #8730) overexpress five FAD mutations: the APP(695) transgene contains the Swedish (K670N, M671L), Florida (I716V), and London (V7171) mutations and the PSEN1 transgene contains the M146L and L286V FAD mutations [30, 32]. Cohort of male and female LOAD1, 5XFAD, and C57BL/6J (B6) controls mice were assayed for serum and brain metabolomics at six months of age (Table 1).

**Table 1.** Number of biological replicates for mouse serum and brain tissue metabolomics. Serum and brain samples were obtained from each mouse.

|  | Female | Male |
| --- | --- | --- |
| 5XFAD | 12 | 13 |
| LOAD1 | 13 | 10 |
| C57BL/6J | 14 | 12 |

### Metabolomics data acquisition and processing

A uniform technical approach was used for both mouse and human metabolomics processing. Full sample preparation and analysis protocols are described on the AD Knowledge Portal (https://adknowledgeportal.synapse.org/Explore/Studies/DetailsPage?Study=syn22313528). In brief, metabolites were measured with the targeted AbsoluteIDQ-p180 metabolomics kit (BIOCRATES Life Science AG, Innsbruck, Austria), with an ultra-performance liquid chromatography tandem mass spectrometry (MS/MS) system (Acquity UPLC (Waters), TQ-S triple quadrupole MS/MS (Waters)), which provides measurements of up to 186 endogenous metabolites. Sample extraction, metabolite measurement, identification, quantification, and primary quality control (QC) followed standard procedures [33]. In serum metabolome, 27 metabolites were removed due to missing values in more than 20% of samples and four metabolites were excluded due to more than 20% coefficient of variation. In brain metabolome, 20 metabolites were removed due to missing values and one metabolite were excluded due to higher coefficient of variation. After median-based batch correction, metabolite concentrations were log2-transformed and missing values for 79 serum metabolites and seven brain metabolites were imputed using kNN (k-nearest-neighbor, k=10) method [34].

### ADNI and ROS/MAP Metabolomics data

Processing of ADNI P180 measurements is described in detail here [12]. In brief, metabolites with >20% missing values were excluded and batch correction was performed using a cross-plate mean normalization procedure using NIST standard plasma metabolite concentrations. Metabolites having a coefficient of variation > 20% or an intra-class correlation < 65% in replicate samples were removed. Missing values were imputed using minimum imputation (set to half of the plate-specific lower limit of detection). Metabolite levels were log2-transformed and multivariate sample outliers were excluded. Finally, metabolites were adjusted for significant medication effects using stepwise backwards selection.

In ROS/MAP, brain and serum metabolites and samples with over 25% missing values were filtered out. Quotient normalization was used to correct for sample-wise variation across the metabolites [35]. Metabolite values were subsequently log2-transformed and the remaining missing values were imputed using kNN as for the mouse data. Data processing was performed using standardized pipelines in the toolbox maplet [36]

### Principal Component Analysis

We analyzed a total of 142 metabolites present in both serum and brain metabolome from 74 samples originating from different mouse models. We extracted the principal components using the singular value decomposition technique for both metabolomes separately and plotted using ggplot2 visualization package in R.

### Association analyses for the mouse metabolomic data

Association analyses of AD risk factors with metabolite levels were conducted using standard linear regression. The stratification variable sex was excluded as a covariate in the sex-stratified analyses. We transformed covariate-adjusted effect sizes to sample size-weighted standardized effects (Cohen’s d). For identifying metabolic sex differences, we used linear regression with metabolite levels as the dependent variable and sex as explanatory variable. To adjust for multiple testing, we used the threshold of Bonferroni significance of 9.09 x 10^-4^ as determined in a recent study [12]. We calculated the power for two-sample t-tests to identify significant sexual dimorphisms for metabolites with the standardized effect sizes observed in the pooled mouse samples at Bonferroni significance in B6, LOAD1, and 5XFAD mouse models. To assess the significance of heterogeneity between strata, we used the methodology of that is similar to the determination of study heterogeneity in inverse-weighted meta-analysis [37]. We further provide a scaled (0–100%) index of percent heterogeneity similar to the I^2^ statistic.

### Association Analysis for the ADNI cohort

Association analysis of the CSF Aβ_1–42_ pathology with blood metabolite levels from the ADNI cohort followed published protocols in Arnold et al. [12] with a few adjustment. We performed the multivariable linear regression analysis using metabolite concentrations as dependent variable to test associations of the CSF Aβ_1–42_ pathology with concentrations of 139 blood metabolites in place of logistic regression approach used by Arnold et al. [12] to be consistent with approach used for mouse metabolomics data for comparison purpose.

### Correlation Analysis

First, we performed the multivariable linear regression analysis using metabolite concentrations of 70 glycerophospholipids (PC species) as dependent variables to determine the effect of APOE4 in serum metabolomics data from mice, ADNI, and ROS/MAP carriers. We also fit a multivariable linear regression model for brain metabolomics data from mouse models and ROS/MAP cohort of the same individuals to measure the effect of APOE4 on glycerophospholipids (PCs) levels. For serum and brain metabolomics data from mouse models, we used 5XFAD genotype and sex as a covariate. For serum metabolomics data from the ADNI cohort, we used pathological CSF Aβ_1–42_, age, sex, cohort, and BMI as covariates. For serum metabolomics data from the ROSMAP cohort, we used pathological amyloid levels, fasting status, age at visit, sex, education and BMI as covariates. For brain metabolomics data from the ROS/MAP cohort, we used pathological amyloid levels, age, sex, education, and BMI as covariates. Finally, we measured the Pearson correlation between effects of *APOE4* on glycerophospholipids (PCs) metabolite levels in human and mouse models. We also measured Pearson correlations between the effects of *APOE4* presence on glycerophospholipids (PCs) metabolite levels in mouse serum and brain metabolomes.

## RESULTS

### Sex is a major separator in both serum and brain metabolome

Principal component analysis indicated serum metabolites separating the samples by sex of the mice along the first principal component (45% of total variance), whereas we observed a slight gradient of discrimination by genotype along the second principal component (13% of total variance) (Figure 1A). Specifically, LOAD1 mice are segregated from B6 and 5XFAD mice. In brain metabolomes, principal component analysis did not reveal any clear separation between groups (sex or genotype). However, a gradient of discrimination of the 5XFAD samples from B6 and LOAD1 samples along the first principal component (46% of total variance), whereas most of the male and female samples were separated along the second principal component (Figure 1B). Overall, this suggests global sex-specific differences in both serum and brain metabolomes.

**Figure 1:**
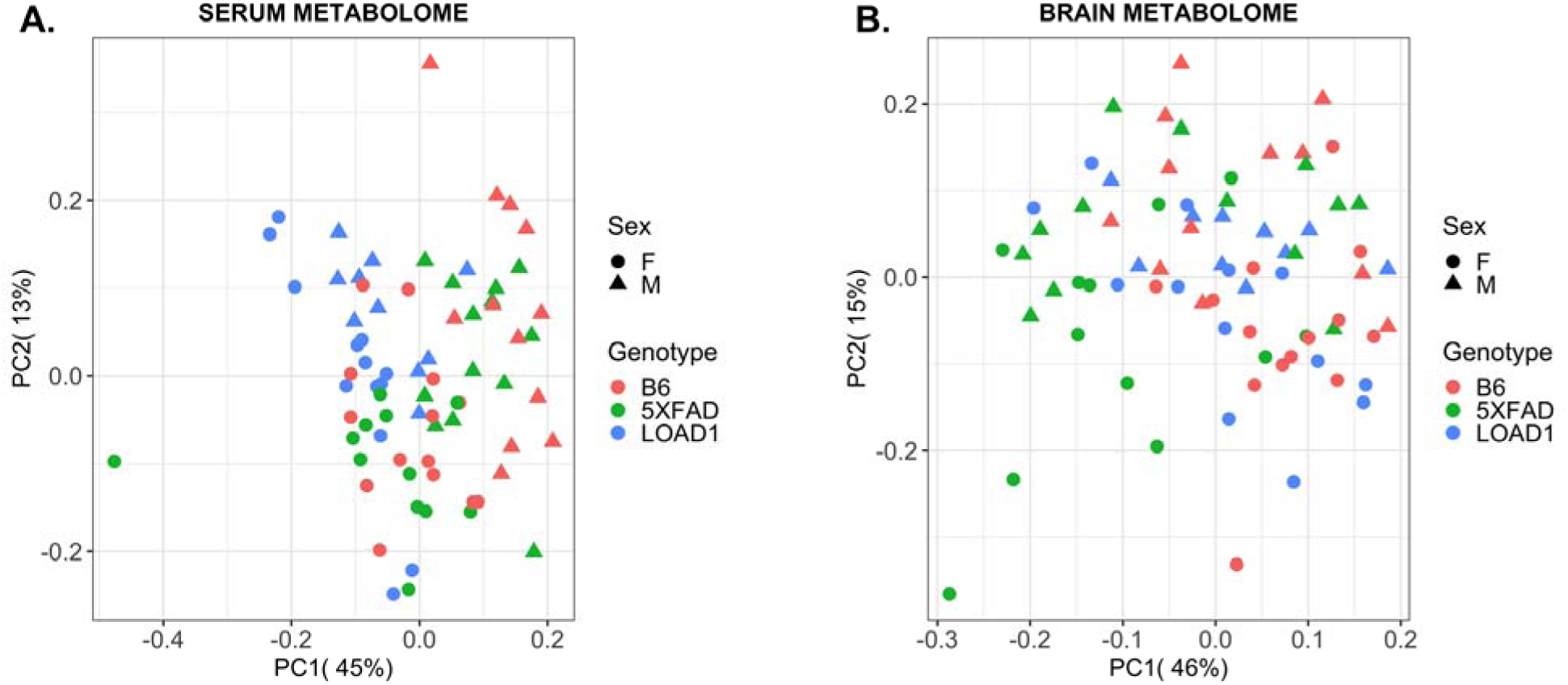
Principal components analysis of serum and brain metabolomes from mouse models of AD. **(A)** In serum metabolome, we observed a slight gradient of discrimination of the LOAD1 samples from B6 and 5XFAD samples along the second principal component; **(B)** In brain metabolome, principal component analysis revealed a gradient of discrimination of the 5XFAD samples from B6 and LOAD1 samples along the first principal component.

### Sex-associated differences significantly differ in AD mouse models

Next, we investigated sex-specific associations of metabolites in all mice together, as well as in each mouse model (B6, LOAD1, and 5XFAD) separately. Then, we tested whether these sex-specific differences are altered in LOAD1 and 5XFAD mice compared to B6 mice to determine strain-specific sex differences.

#### Mouse serum metabolome

In the complete cohort (N = 74), we identified 73 out of 142 metabolites significantly associated with sex after multiple testing correction (p < 9.09 x 10^-4^) while adjusting for genotypes (Figure 2A, Supplementary Table 1). Fifty-six of these metabolites had higher levels in males, and 17 metabolites had higher levels in females. The majority of glycerophospholipids (PCs) were more abundant in male mice, while levels of few amino acids (alanine, isoleucine, serine, and threonine) and majority of sphingolipids were more abundant in female mice (Supplementary Table 1). Notably, 54 of these sex-specific associations were also observed in a recent study from the ADNI cohort [12]. Further, stratification by genotypes revealed that 13 of the 73 metabolites showing significant sex differences in the complete cohort were also significant in each of the three genotypes (B6, LOAD1, and 5XFAD) alone, whereas 14 metabolites showed no significant difference in any genotype (Supplementary Table 1, Figure 2A). Significant sex differences limited to one genotype were found for three metabolites (lysoPC a C20:4, SM (OH) C24:1, ADMA) in LOAD1 mice, for two metabolites (C3-DC (C4-OH), PC ae C30:1) in the 5XFAD mice, and for 21 metabolites in B6 mice. Every significant sex difference found in each genotype alone was also significant in full cohort except one; PC aa C38:4 showed a significant sex difference in only LOAD1 genotype with higher levels in females (Supplementary Table 1).

**Figure 2:**
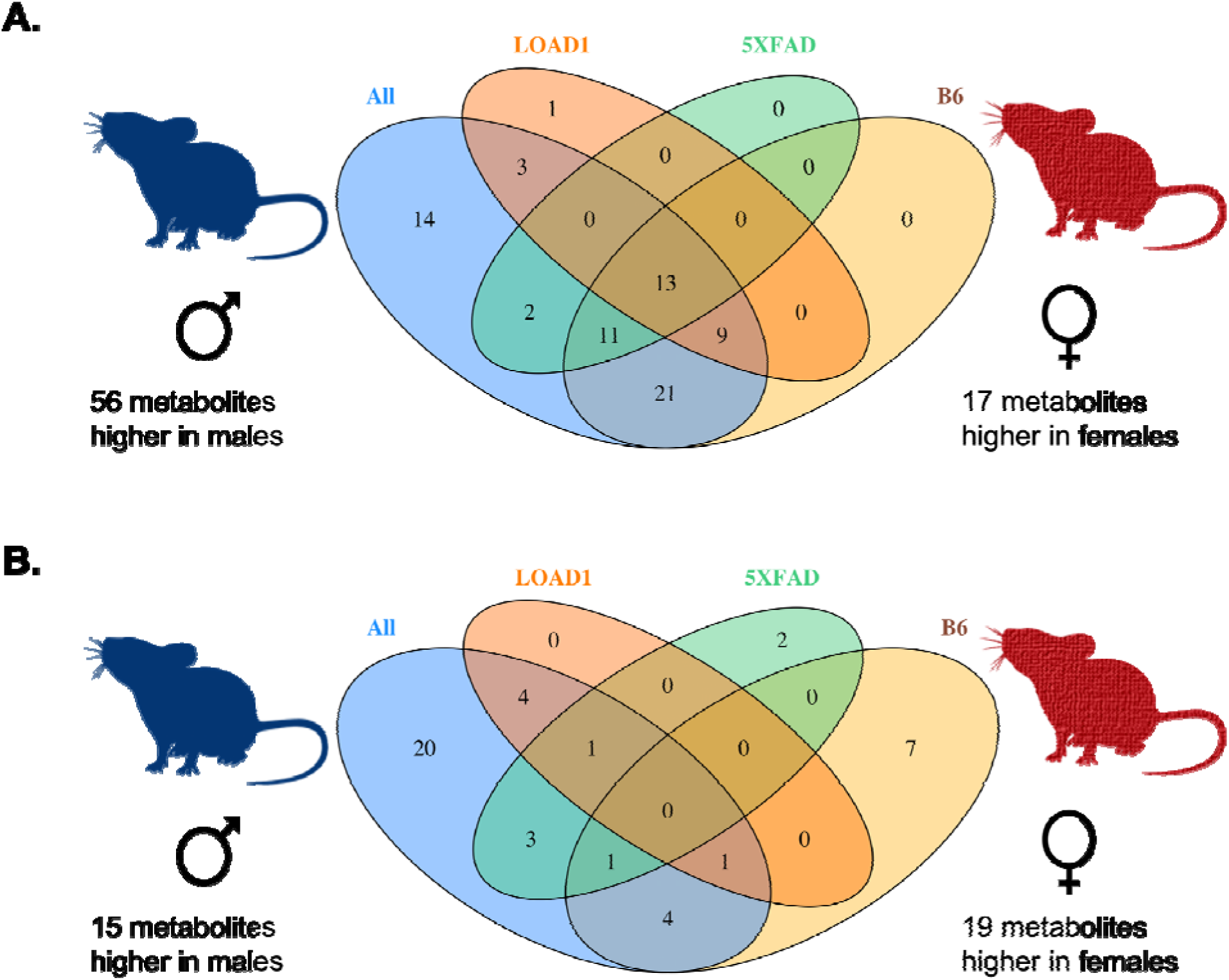
Sex-metabolite associations in mouse models. We measured metabolites significantly associated with sex in the full mouse cohort and in each genotype separately. **(A)** Number of significantly altered metabolites in serum metabolome. **(B)** Number of significantly altered metabolites in brain metabolome. The Venn diagrams show the overlap between metabolites associated with sex in full cohort and in each genotype for both serum and brain metabolomes.

Further, comparisons of beta estimates for sex between 5XFAD and B6 groups showed significant effect heterogeneity (P_HET_ < 0.05) for five metabolites (C18:1, C18:2, PC aa C34:2, PC aa C36:2, SDMA). All five metabolites had a change in direction of abundance between sex in the 5XFAD mice compared to B6 mice. Notably, none of these metabolites showed significant sex specific differences either in combined mouse samples or by individual genotype. Similarly, comparisons of beta estimates for sex between LOAD1 and B6 groups showed significant effect heterogeneity (P_HET_ < 0.05) for 31 metabolites, out of which 23 were glycerophospholipids and three were amino acids (valine, arginine, and phenylalanine) (Supplementary Table 2). Most of these metabolites exhibited a change in direction of abundance between sex in LOAD1 compared to B6 controls. These observations indicated that sex differences of serum metabolite levels were significantly affected in late-onset AD model compared to control mice.

#### Mouse brain metabolome

Next, we investigated sex-specific associations of metabolites and sex-associated differences in brain metabolomes of mouse models. In the complete cohort (N = 74), we identified 34 out of 142 metabolites significantly associated with sex after multiple testing correction (p < 9.09 x 10^-4^) when adjusting for genotypes (Figure 2B, Supplementary Table 1). A total of 15 metabolites had higher levels in males including ten glycerophospholipids, four sphingolipids (SMs), and one biogenic amines, while 19 metabolites had higher levels in females including nine glycerophospholipids, four amino acids, and six acylcarnitines (Supplementary Table 1). Stratification by genotypes revealed 14 of the 34 metabolites showing significant sex differences were significant in at least one of the genotypes (B6, LOAD1, and 5XFAD), whereas 20 showed no significant sex difference in any genotype. Significant sex differences limited to one genotype were found for four metabolites (C18:1, PC aa C32:3, PC ae C36:4, t4-OH-Pro) in LOAD1 mice, for three metabolites (C18, PC aa C40:4, PC ae C36:5) in the 5XFAD mice, and for four metabolites (PC ae C42:2, SM C24:0, SM (OH) C22:1, SM (OH) C24:1) in the B6 mice. Significant sex differences for two metabolites (Spermidine, Spermine) in 5XFAD and seven metabolites (PC ae C30:0, PC ae C30:1, PC ae C30:2, PC ae C34:0, PC ae C36:0, SM C16:0, SM C18:0) in B6 were not significant in full cohort (Supplementary Table 1, Figure 2B).

Comparisons of beta estimates for sex between 5XFAD and B6 groups showed significant effect heterogeneity (P_HET_ < 0.05) for 29 metabolites, which include 21 PCs (majority of which were ester-containing PCs), five SMs, and three biogenic amines (Supplementary Table 3). All but two metabolites showed a change in direction of abundance between sex (higher levels in females compared to males) in the 5XFAD mice compared to B6 mice. Similarly, comparisons of beta estimates for sex between LOAD1 and B6 groups showed significant effect heterogeneity (P_HET_ < 0.05) for 13 metabolites, that included nine glycerophospholipids (the majority of which were ether-containing PCs), two long chain acylcartines (C16:1, C18:2), one biogenic amine (SDMA), and one hydroxy-SM (SM (OH) C22:1) (Supplementary Table 3). All but three metabolites exhibited a change in direction of abundance between sex (higher levels in females compared to males) in LOAD1 mice compared to B6 mice. Moreover, three metabolites (PC aa C38:6, PC ae C32:1, PC ae C44:6) showed significant heterogeneity in both 5XFAD and LOAD1 mouse models compared with controls. In summary, we found that sex differences of brain metabolite levels varied by genotype in mouse models, with more pronounced effects in 5XFAD.

Moreover, we identified 19 metabolites showed sex-specific differences in both serum and brain metabolomes in the combined analysis of all mice. Notably, none of the metabolites showed sex differences in both serum and brain metabolomes of the LOAD1 mice, while only one metabolite PC aa C40:4 showed sex differences in both serum and brain metabolomes of the 5XFAD mice but levels of PC aa C40:4 were higher in males and females in serum and brain metabolomes, respectively.

### Sex stratified associations of metabolites with AD risk factors

Next, we investigated associations of metabolites with AD risk factors and whether sex modifies the associations between AD risk factors and metabolites concentrations. We tested for associations of the 5XFAD and the LOAD1 genotypes with concentrations of 142 metabolites in both serum and brain metabolomes from same 74 individual animals. We did this in the full cohort and separately in each sex using multivariable linear regression, followed by analysis of heterogeneity of effects between sexes. All metabolite-genotype associations were deemed significant that fulfilled at least one of the three criteria as described in Arnold et al. 2020 [12]: (i) associations Bonferroni significant (at a threshold of p < 9.09 × 10^−4^) in the full cohort; (ii) associations Bonferroni significant in one sex; (iii) associations that showed suggestive significance (p < 0.05) in one sex coupled with significance for effect heterogeneity between female and male effect estimates. These significant associations were further classified into homogeneous (metabolites with similar effects in their association to the risk factors for both sexes), heterogeneous (metabolites with different effects in both sexes leading to significant heterogeneity) and sex-specific effects (effects that were Bonferroni significant in only one sex with significant effect heterogeneity between males and females). We compared our findings in mouse models with recent results from the ADNI cohort [12], and therefore examined the association of metabolites with each genotype for our mice cohort using similar methods.

#### Mouse serum metabolome

In mouse in serum metabolome, we identified 83 metabolites significantly associated with LOAD1 genotype, while only seven metabolites were significantly associated with 5XFAD genotype (Supplementary Table 4, Figure 3A-B). Among the LOAD1 genotype associated metabolites, we found 32 metabolites (24 PCs and 8 SMs) with homogeneous associations. Next, we identified seven associations with heterogeneous effects: one acylcarnitine (C16-OH), one biogenic amine (alpha.AAA), and four amino acids (Ile, Phe, Tyr, Val) have larger effect size in males, whereas ADMA showed stronger associations with females (all I^2^ >50%). Further, male-specific effects were seen for 36 metabolites associated with LOAD1, which included 32 PCs, three SMs, and one acylcarnitines (C14:1-OH). Overall, serum levels of these significant metabolites were reduced in LOAD1 mice compared to B6 controls, except for the four amino acids (Ile, Phe, Tyr, Val) and one biogenic amine (alpha.AAA).

**Figure 3:**
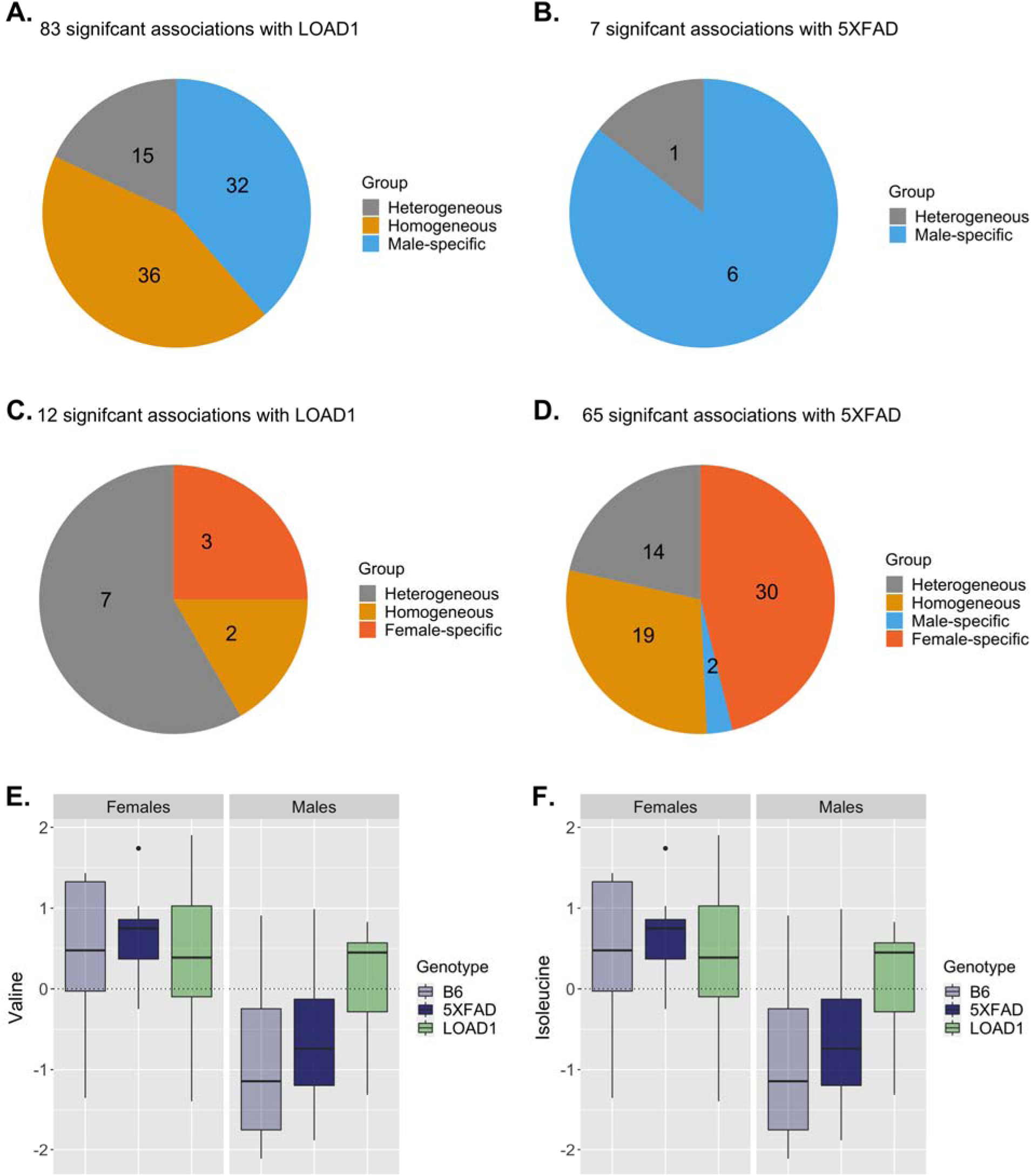
Metabolite associations with LOAD1 and 5XFAD genotype stratified by sex in mouse models. In the serum metabolome, **(A)** 83 metabolites were significantly associated with LOAD1 and **(B)** seven metabolites were significantly associated with 5XFAD. **(C)** In the brain metabolome, 12 metabolites were significantly associated with LOAD1 and **(D)** 65 metabolites were significantly associated with 5XFAD. **(E-F)** Valine and Isoleucine levels in B6 control, 5XFAD, and LOAD1 male and female mice. Levels of these amino acids in serum metabolome are significantly higher (p<0.05) in male LOAD1 compared to male controls.

For 5XFAD, we found five metabolites with heterogenous effects: acylcarnitines (C18:1, and C18:2) showed stronger associations with females, whereas PC aa C34:2, PC aa C36:2 and one biogenic amine (SDMA) yielded stronger associations with males (all I^2^ >50%). One male-specific effect was seen for PC ae C38:1 with significant negative association with 5XFAD.

Another metabolite (PC aa C32:1) met the Bonferroni threshold for male but failed to meet threshold for sex effect heterogeneity. Serum levels of these 5XFAD associated metabolites were also reduced in 5XFAD mice compared to B6 controls, except for one biogenic amine (SDMA). Further, we did not observe female-specific effects for any metabolite associated with either genotype.

#### Mouse brain metabolome

In the mouse brain metabolome, we identified 12 metabolites significantly associated with LOAD1, while 65 metabolites were significantly associated with the 5XFAD genotype (Supplementary Table 4 Figure 3C-D). Out of 12 metabolites associated with LOAD1, two had homogenous effects (PC aa C42:4, PC ae C34:3) with strong positive associations. Next, we identified 6 associations with heterogeneous effects: acylcarnitine (C18:2), and biogenic amine (SDMA), with larger effect size in males, whereas PCs (PC aa C28:1, PC ae C40:1, PC ae C44:6), and one-hydroxy sphingolipids (SM (OH) C22:1) showed stronger positive associations with females (all I^2^ >50%). Further, female-specific effects were seen for three ether-containing PCs (PC ae C32:1, PC ae C34:0, PC ae C36:0), with significant positive correlations with the LOAD1 genotype. Levels of these significant metabolites with homogenous and female-specific were increased in LOAD1 mice compared to B6 controls, while levels of metabolites with heterogenous effects were reduced in males and increased in females. Overall, we observed increased levels of these metabolites in female LOAD1 mice compared to female B6.

Out of 65 metabolites associated with 5XFAD genotype, we found 19 metabolites with homogenous effects, that includes 15 PCs, one SM (SM C16:1), one amino acid (lysine), and two biogenic amines (creatinine and t4-OH-Pro). Six PCs and one hydroxy sphingolipids (SM (OH) C22:2) showed heterogenous effects with stronger positive associations with females (I^2^ >50%). We also identified 30 metabolites with female-specific effects: 21 PCs, 7 SMs (including 4 hydroxy SMs), one biogenic amine (putrescine), and one acylcarnitine (C18). Levels of these 5XFAD associated metabolites were higher in 5XFAD mice compared to B6 controls. Male-specific effects were also seen for two biogenic amines (spermidine, spermine), with strong negative associations with 5XFAD.

Upon comparing these brain associations with associations in the serum metabolome, we observed that out of 65 metabolites associated with 5XFAD in brain metabolome, only three metabolites were associated with serum in 5XFAD, while 43 were associated with serum in LOAD1. This suggests that AD-related effects might be present in different tissues in the different mouse models. Similarly, out of 12 metabolites associated with LOAD1 in the brain, ten were associated with LOAD1 and two were associated with 5XFAD in serum. Metabolites commonly associated with either genotype across serum and brain metabolomes were generally glycerophospholipids and sphingolipids. Some amino acids such as valine and isoleucine were significantly associated with the LOAD1 genotype in serum metabolome with larger effect size in males (Figure 3E-F, Supplementary Table 3), while two amino acids (lysine and arginine) and five biogenic amines (spermidine, spermine, creatinine, putrescine and t4-OH-Pro) were specifically significantly associated with 5XFAD in brain metabolome. Notably, levels of the significantly associated metabolites were higher in brain metabolome and reduced in serum metabolome compared to respective controls. We also compared the effect of LOAD1 on glycerophospholipids (the largest class of metabolites in panel) levels between serum and brain metabolome and observed a significant negative correlation (r = -0.31,p = 0.01) between them (Figure 5A).

### Comparison of mouse metabolome profile with human metabolome study

Next, we compared our results with human metabolomic profiles from two independent cohorts: (i) the Alzheimer’s Disease Neuroimaging Initiative (ADNI) cohort, for which we have serum metabolomics data from 1517 participants [12]; and (ii) the Rush Religious Order Study and Memory and Aging Project (ROS/MAP) [31], for which we had access of p180 metabolites data from both serum and brain from 92 participants.

### The ADNI Cohort

We investigated whether serum metabolites reported to be significantly associated with AD biomarkers in the ADNI cohorts [12] were also associated with AD risk factors in mouse serum metabolome. Arnold et al. [12] have used multivariable logistic regression to measure associations of pathological Aβ_1–42_ with metabolite concentrations as explanatory variable. For uniformity with our mouse data analysis, we re-assessed the association analysis for the ADNI human serum metabolomics data using multivariable linear regression approach using metabolite concentrations as dependent variable to test associations of the CSF Aβ_1–42_ pathology with concentrations of 139 serum metabolites. We were able to recover same metabolites significantly associated with CSF Aβ_1–42_ pathology as reported in the study [12].

Interestingly, we observed that four metabolites significantly associated with CSF Aβ_1–42_pathology (PC ae C44:4, PC ae C44:5, PC ae C44:6, and Valine) were also significantly associated with the LOAD1 genotype (Supplementary Table 3). Three out of these four metabolites (PC ae C44:4, PC ae C44:5, and PC ae C44:6) also showed Bonferroni-significant associations with pathological CSF Aβ_1–42_ in APOE ε4 carriers [12]. Further, we noticed a similar decrease in these metabolites levels in serum of APOE ε4 carriers (without CSF Aβ_1–42_ pathology), as we have seen in LOAD1 mice that carry the ε4 variant but do not exhibit amyloid plaques (Figure 4). However, none of the significant associations reported for pathological CSF Aβ_1–42_ were significantly associated with the 5XFAD genotype in mouse serum metabolome.

**Figure 4:**
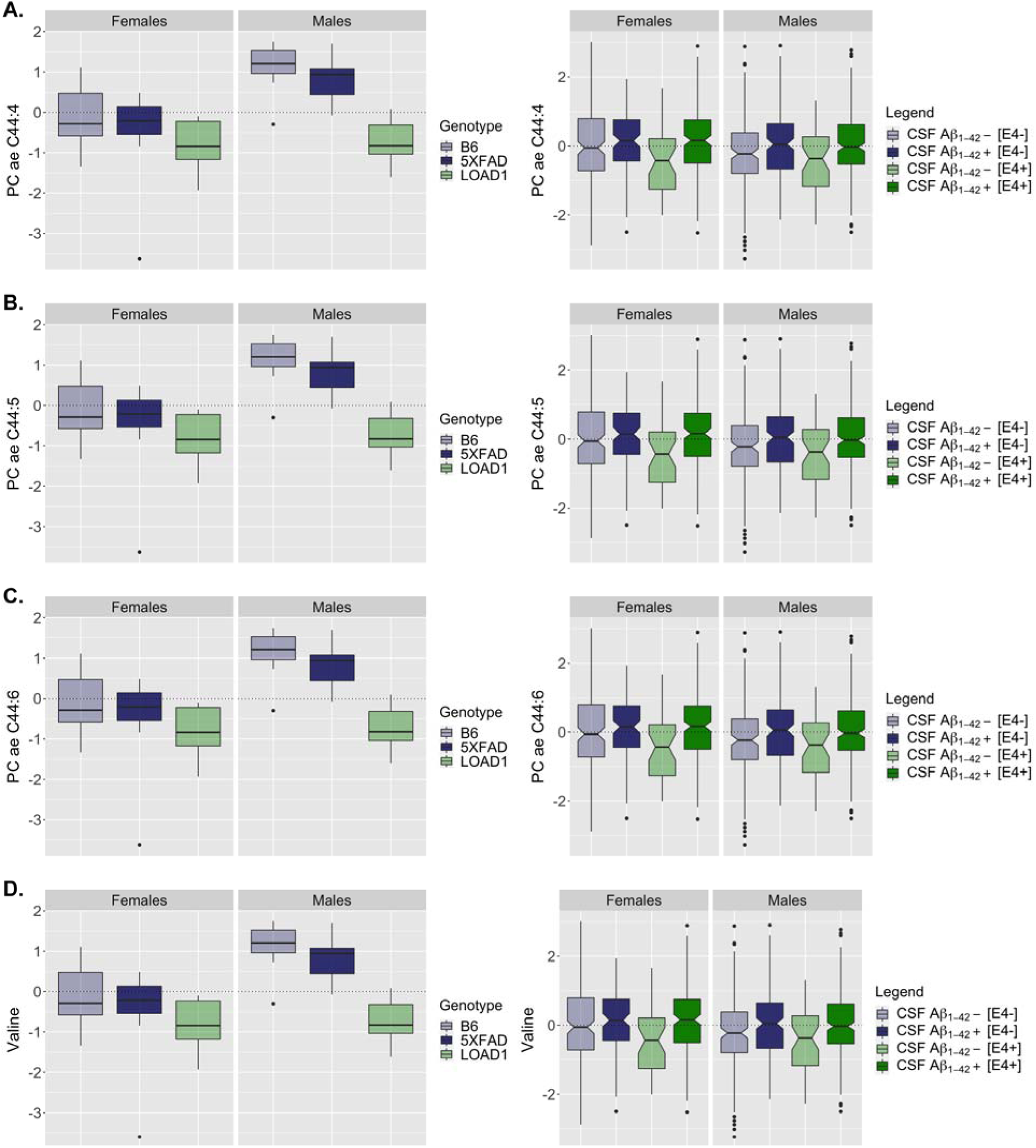
Levels of metabolites significantly associated with CSF Aβ_1–42_ pathology in humans and LOAD1 genotype in mouse models. Levels of **(A)** PC ae C44:4; **(B)** PC ae C44:5; **(C)** PC ae C44:6; and **(D)** Valine; in human subjects (right panels) and mouse models (left panels) separately for both sexes. In humans, subjects were stratified by *APOE* ε4 carriers as well CSF Aβ_1–42_pathology.

Further, we measured correlation between effects of APOE ε4 on glycerophospholipids levels in serum metabolomics data from the ADNI cohort and mouse models. We identified significant positive correlation (r = 0.31, p = 0.008) between effects of APOE ε4 on glycerophospholipids levels in human and mouse serum, suggesting similar decrease in the glycerophospholipids in serum of APOE ε4 carriers compared to non-carriers and indicates serum-based effect of APOE ε4 genotype (Figure 5B).

**Figure 5:**
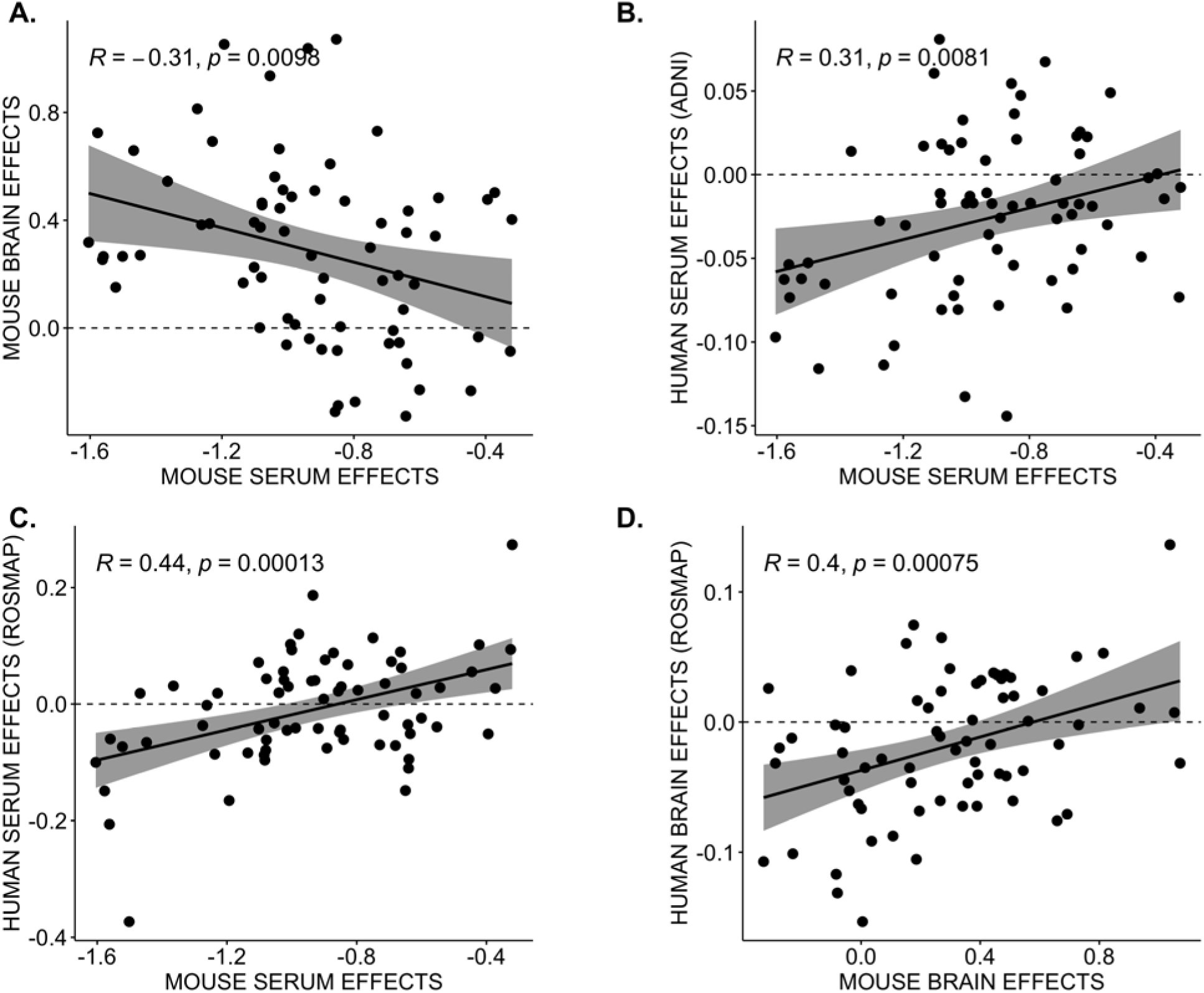
Correlation between human and mouse metabolomic signatures in *APOE* ε4 carriers. *APOE* ε4 effects on glycerophospholipids levels were measured between mouse and human metabolomes with Pearson correlation (*R*) and statistical significance. Each point is an effect estimate for a single glycerophospholipid species. **(A)** Mouse serum and brain effects showing significant negative correlation in LOAD1 mice. **(B)** ADNI human and mouse serum effects showing significant positive correlation. **(C)** ROS/MAP human and mouse serum effects showing significant positive correlation. **(D)** ROS/MAP human and mouse brain effects showing significant positive correlation.

### ROS/MAP Cohort

Next, we carried out multivariable linear regression analysis on ROS/MAP serum and brain metabolomics data using metabolite concentrations of glycerophospholipids as dependent variables to measure the effects of the APOE ε4 genotype versus other *APOE* genotypes. We then measured the correlation between the effects of APOE ε4 on glycerophospholipids levels in serum metabolomics data from ROS/MAP cohort and *APOE4* carriers among mouse models (*i.e.* LOAD1 mice). We observed a significant positive correlation (r = 0.44, p = 0.0001) between the effects of APOE ε4 on glycerophospholipids levels in human and mouse serum, suggesting a similar decrease in the glycerophospholipids in serum of APOE ε4 carriers compared to non-carriers and indicates the serum-based effect of APOE genotype (Figure 5C). We also measured the correlation between effects of APOE ε4 on glycerophospholipids levels in brain metabolomics data from ROS/MAP cohort and mouse models and found a significant positive correlation (r = 0.40, p = 0.0008) (Figure 5D), suggesting a simultaneous increase in the same metabolites in the brain, consistent across humans and mice. We also compared effect of APOE ε4 on glycerophospholipids levels in human serum and brain metabolome from the ROS/MAP cohort, but we did not observe a significant negative correlation (r = 0.11,p = 0.4) as observed in mouse models. In summary, we observed the same effect directions in humans and mice from both serum and brain metabolites when compared separately, and human serum-to-brain comparisons may suffer from limited power.

## DISCUSSION

### Summary

In this study, we have systematically investigated alternations in abundances of 142 metabolites in the serum and brain metabolome of the 5XFAD amyloid mouse model and a more recently created LOAD1 (APOE4.Trem2^R47H^) mouse model with late-onset AD genetics. We assessed the sex differences for metabolic associations in each mouse model and investigated changes in sexual dimorphisms of metabolic levels when compared to B6 control mice. A complex pattern of sex and genotype effects were observed, with the most significant effects occurring in the brain of 5XFAD mice and serum of LOAD1 mice. In order to assess the translational relevance of the mouse models, we compared our findings with recent metabolomic studies from two human cohorts [12, 31]. We observed similar glycerophospholipid signatures in human and mouse *APOE* ε4 carriers in both brain and serum.

### The two mouse strains exhibit distinct sex-specific biology, both with potential relevance to dementia

Overall, we observed distinct sex differences for the two strains. Significant sexual dimorphism of serum metabolic levels were observed in mice carrying the *APOE* ε4 variant. In the serum metabolome, we observed that most of glycerophospholipids were significantly associated with males and showed elevated levels compared to females, while few sphingomyelins and amino acids were more abundant in females. Multiple serum metabolites with significant higher levels in B6 (controls) females (specifically amino acids including valine and arginine) showed reduced levels in female mice carrying *APOE* ε4 variant (LOAD1) compared to their male counterparts, while other metabolites such as phospholipids and some long chain acylcarnitines showed reduced levels in male mice carrying *APOE* ε4 variant compared to females. Brain metabolomes showed more significant sexual dimorphisms in 5XFAD mice and brain metabolites such as some biogenic amines, phospholipids, and sphingomyelins that had greater abundance in B6 males showed higher levels in the 5XFAD females. This suggests that metabolic sex differences changed owing to presence of *APOE* ε4 and 5XFAD variant. However, few amino acids such as alanine, serine, threonine and tryptophan exhibited significant increased levels in both brain and serum metabolome of female mice compared to male mice in all three genotypes, while some phospholipids were significantly elevated in brain and serum metabolome of male mice compared to female mice all three genotypes. These results suggest that even though both mouse models are intended for use in Alzheimer’s disease research, the two strains have distinct sex-specific biology with potential relevance to dementia.

### The LOAD1 mouse model is appropriate for the study of AD metabolite biology

Our analysis of both brain and serum metabolomes from each animal suggests distinct tissues of action for the two genetic constructs. Sex-stratified metabolite associations with genotypes identified that serum metabolites were more significantly associated with LOAD1 genotype, while brain metabolites were more significantly associated with the 5XFAD genotype. Notably, we identified a heterogenous association of valine in the LOAD1 genotype, with reduced levels in female mice but increased levels in males. Studies have associated the reduced level of valine in serum with cognitive decline and brain atrophy in AD [38], and also suggested valine as a marker for increased female vulnerability to AD [12, 38]. Interestingly, we observed increased levels of biogenic amine putrescine, specifically in the brain of female 5XFAD. As significant increased level of putrescine have been reported in brain tissue from AD patients [39], Similar elevation were also observed in the APP/PS1 mice at 6 months [26], suggesting Aβ causes up-regulation of polyamine uptake and increased ornithine decarboxylase activity, which leads to increased polyamine synthesis [40, 41], which in return causes dysfunction of the NMDA receptor leading to the neuronal excitotoxicity which occurs in AD [42]. Two other biogenic amines (spermidine, spermine) also exhibit increased but insignificant level in the 5XFAD female brain. Pan et al., [26] reported that putrescine precedes both spermidine and spermine in the biochemical pathway and observed significant increase in both spermidine and spermine in female brain at later time point. This suggests that our relatively young mice (6 months of age) would exhibit more significant changes in these biogenic amines at later ages. The metabolite Asymmetric dimethylarginine (ADMA), which is an endogenous inhibitor of nitric oxide synthase, has been found to be higher in plasma from AD patients [43]. Inhibition of endothelial nitric oxide synthesis by ADMA impairs cerebral blood flow, which may contribute to the development of Alzheimer’s disease [43]. We also observed a significant positive association of ADMA with female LOAD1 in serum metabolome, suggesting that the LOAD1 mouse model is appropriate for the study of relevant AD biology.

### 5XFAD amyloidogenic mouse is a relevant model for acylcarnitine alterations in AD

We identified either female-specific or heterogeneous associations of the sphingomyelins with higher levels in the brain metabolome of 5XFAD mice. Sphingomyelins are precursors for ceramide production and their accumulation suggested to induce apoptosis and seems to worsen neurodegeneration by increasing amyloid beta biosynthesis and promoting gamma-secretase processing of amyloid precursor protein [44-46]. Higher levels of sphingomyelins and glycerophospholipids in the six-months-old mouse brains are indicative of early neurodegeneration and loss of membrane functions. Acylcarnitines have important functions in the brain such as mitochondrial function, energetics, and neurotransmission and have been linked with AD related pathology [12, 47, 48]. We observed a significant female specific association of higher levels of acylcarnitine C18 with the 5XFAD, suggesting sex-specific accumulation of long chain fatty acids in females. Increased level of C18 has been also previously reported in mild cognitive impaired patients with CSF Aβ_1–42_ pathology [47]. In contrast to ADMA in LOAD1, these findings suggest the 5XFAD amyloidogenic mouse as a relevant model for acylcarnitine alterations in AD.

### Serum biomarkers are informative, but their effects in the brain cannot be directly extrapolated

Glycerophospholipids (PCs and LysoPCs) are the major class of complex lipids playing essential roles in neural membrane formation and intraneuronal signal transduction [49]. We identified that serum levels of glycerophospholipids were reduced in LOAD1 mice compared to B6 controls, while levels of these metabolites in the same animals were greater in both male and female brains of LOAD1 and 5XFAD mice. We also compared the LOAD1 genotype effects on glycerophospholipid levels between serum and brain metabolome and observed a significant negative correlation. This suggests a correspondence between brain and serum glycerophospholipid levels, but a negative rather than positive correlation. Similar pattern of contrast concentration level of glycerophospholipids in brain and blood metabolome were also observed in APP/PS1 mouse [26]. These finding imply that although the serum biomarker is informative, effects in the brain cannot be directly extrapolated.

### The translational utility of mouse models for metabolomic studies of AD

We observed the translational relevance for these results in two human AD studies. Our observed effect of LOAD1 on serum glycerophospholipids levels was significantly correlated with serum effects of the *APOE* ε4 variant in ADNI, showing a similar decrease in the same metabolites in serum of *APOE* ε4 carriers compared to non-carriers and indicates a serum-based effect of *APOE* genotype. We also identified Bonferroni significant associations of PC ae C44:4, PC ae C44:5, and PC ae C44:6 with *APOE* ε4 genotype in mouse models as in the ADNI cohort with consistent effect directions. While all three PCs showed homozygous associations in the ADNI cohort, in mice only PC ae C44:5 showed a homogeneous effect, while associations of PC ae C44:4 and PC ae C44:6 were male-specific. We also confirmed tissue-specific effects of the LOAD1 genotype on metabolite levels for *APOE* ε4 carriers in the ROS/MAP cohort, finding significant correlations between carriers and LOAD1 mice in both brain and serum metabolome. Altogether, this suggests that we were able to identify similar effects of the *APOE* ε4 variant in mouse models as observed on human AD carriers. However, we did not observe consistent sex differences between mice and humans. Contrary to the mouse models, PCs were higher in female humans and amino acids were (with few exceptions like serine and glycine) higher in males [12]. Despite these dissimilarities of some sex differences, we still observed an overlap in associations with mouse genotype and the related human phenotypes, which indicates that the sex differences do not mask associations on the phenotypic/genotypic level. This suggests that, while there is a difference in the metabolic patterns for sex in mice and humans, there is translational utility in mouse models for metabolomic studies of AD.

## Conclusion

In conclusion, metabolomic signatures were notably different between brain and serum in two mouse models of AD at six months of age. The early-onset 5XFAD mice exhibited stronger effects in brain, whereas the late-onset LOAD1 mouse showed more pronounced effects in serum. These findings are consistent with the high levels of neuropathology in 5XFAD mouse brains and the modifications of serum biomarkers in LOAD1 mice. We were able to identify patterns of metabolic changes related to human AD risk in our mice models. These findings strengthen the utilization of mouse models for studying metabolic changes that occur in human AD at early stages. However, there are some limitations in this study. First, we have evaluated metabolomic data from only six-month-old mice, which did not allow to study progressive metabolic changes associated with AD. Studies using transgenic APP/PS1 mice observed that metabolic changes associated with AD pathology appeared first in the brain and later in blood [26]. Therefore, it will be interesting to investigate metabolomics profiles of the LOAD1 and other late-onset mouse models over different extended ages to study progressive metabolic changes associated with AD pathology. Our findings suggest such aging studies can model the different phases of metabolic disturbances which may occur in human AD.

## Data Availability Statement

Metabolomics datasets are available via the AD Knowledge Portal (https://adknowledgeportal.org). The AD Knowledge Portal is a platform for accessing data, analyses, and tools generated by the Accelerating Medicines Partnership (AMP-AD) Target Discovery Program and other National Institute on Aging (NIA)-supported programs to enable open-science practices and accelerate translational learning. The data, analyses and tools are shared early in the research cycle without a publication embargo on secondary use. Data is available for general research use according to the following requirements for data access and data attribution (https://adknowledgeportal.org/DataAccess/Instructions).

For access to content described in this manuscript see:

Jax.IU.Pitt Metabolomics (p180): https://www.synapse.org/#!Synapse:syn22313586

ADNI p180: https://www.synapse.org/#!Synapse:syn7440346

ROS/MAP p180: https://www.synapse.org/#!Synapse:syn26007829

## Acknowledgements and Funding Sources

The results published here are in whole or in part based on data obtained from the AD Knowledge Portal ( https://adknowledgeportal.org). The IU/JAX/PITT MODEL-AD Center was established with funding from The National Institute on Aging (U54 AG054345). Metabolomics data is provided by the Alzheimer’s Disease Metabolomics Consortium (ADMC) and funded wholly or in part by the following grants and supplements thereto: NIA R01AG046171, RF1AG051550, RF1AG057452, R01AG059093, RF1AG058942, U01AG061359, U19AG063744 and FNIH: #DAOU16AMPA awarded to Dr. Kaddurah-Daouk at Duke University in partnership with a large number of academic institutions. As such, the investigators within the ADMC, not listed specifically in this publication’s author’s list, provided data along with its pre-processing and prepared it for analysis, but did not participate in analysis or writing of this manuscript. A complete listing of ADMC investigators can be found at: https://sites.duke.edu/adnimetab/team/. Data was generated by the Duke Metabolomics and Proteomics Shared Resource, a member of the ADMC, using protocols published previously for blood samples (Toledo et al., https://doi.org/10.1016/j.jalz.2017.01.020; St. John-Williams et al., https://doi.org/10.1038/sdata.2017.140; Arnold et al., https://doi.org/10.1101/585455). MA, RKD, JK and RB are additionally supported by the National Institute of Aging of the National Institutes of Health under awards NIA 1U19AG063744 and R01AG069901-01.

**ADNI:** Data collection and sharing for this project was funded by the Alzheimer’s Disease Neuroimaging Initiative (ADNI) (National Institutes of Health Grant U01 AG024904) and DOD ADNI (Department of Defense award number W81XWH-12-2-0012). ADNI is funded by the National Institute on Aging, the National Institute of Biomedical Imaging and Bioengineering, and through generous contributions from the following: AbbVie, Alzheimer’s Association; Alzheimer’s Drug Discovery Foundation; Araclon Biotech; BioClinica, Inc.; Biogen; Bristol-Myers Squibb Company; CereSpir, Inc.; Cogstate; Eisai Inc.; Elan Pharmaceuticals, Inc.; Eli Lilly and Company; EuroImmun; F. Hoffmann-La Roche Ltd and its affiliated company Genentech, Inc.; Fujirebio; GE Healthcare; IXICO Ltd.; Janssen Alzheimer Immunotherapy Research & Development, LLC.; Johnson & Johnson Pharmaceutical Research & Development LLC.; Lumosity; Lundbeck; Merck & Co., Inc.; Meso Scale Diagnostics, LLC.; NeuroRx Research; Neurotrack Technologies; Novartis Pharmaceuticals Corporation; Pfizer Inc.; Piramal Imaging; Servier; Takeda Pharmaceutical Company; and Transition Therapeutics. The Canadian Institutes of Health Research is providing funds to support ADNI clinical sites in Canada. Private sector contributions are facilitated by the Foundation for the National Institutes of Health (www.fnih.org). The grantee organization is the Northern California Institute for Research and Education, and the study is coordinated by the Alzheimer’s Therapeutic Research Institute at the University of Southern California. ADNI data are disseminated by the Laboratory for Neuro Imaging at the University of Southern California.

**ROS/MAP:** Study data provided through NIA grant 3R01AG046171-02S2 was awarded to Rima Kaddurah-Daouk at Duke University, based on specimens provided by the Rush Alzheimer’s Disease Center, Rush University Medical Center, Chicago. Data collection was supported through funding by NIA grants P30AG10161 (ROS), R01AG15819 (ROSMAP; genomics and RNAseq), R01AG17917 (MAP), R01AG30146, R01AG36042 (5hC methylation, ATACseq), RC2AG036547 (H3K9Ac), R01AG36836 (RNAseq), R01AG48015 (monocyte RNAseq) RF1AG57473 (single nucleus RNAseq), U01AG32984 (genomic and whole exome sequencing), U01AG46152 (ROSMAP AMP-AD, targeted proteomics), U01AG46161(TMT proteomics), U01AG61356 (whole genome sequencing, targeted proteomics, ROSMAP AMP-AD), the Illinois Department of Public Health (ROSMAP), and the Translational Genomics Research Institute (genomic). Additional phenotypic data can be requested at www.radc.rush.edu.

